# Reduced Myonuclear Number Is Accompanied by More Regular Longitudinal Nuclear Positioning During Muscle Growth

**DOI:** 10.64898/2026.05.05.722961

**Authors:** Kenth-Arne Hansson, Mikkel Elle Lepperød

## Abstract

Skeletal muscle fibers are large syncytial cells containing hundreds of spatially distributed myonuclei that collectively provide transcriptional output across a shared cytoplasm. Whether the spatial organization of myonuclei adjusts when myonuclear accretion does not keep pace with cytoplasmic growth remains unresolved. We therefore asked whether a smaller myonuclear population is distributed more uniformly within a growing fiber, potentially limiting cytoplasmic regions disproportionately remote from a source of nucleus-derived products. To test this hypothesis, we analyzed three-dimensional myonuclear organization throughout postnatal growth in a mouse model in which inducible Myomaker deletion in muscle stem cells restricts their fusion with growing fibers and thereby limits the addition of new myonuclei. Across 1,029 EDL fibers containing nearly 15,000 myonuclei, sampled from postnatal day 13 (P13) to approximately 5 months of age (P150), the resulting reduction in myonuclear number left fewer nuclear sites of transcriptional output within the growing fiber and increased the characteristic distance between neighboring myonuclei. Despite this greater separation, the myonuclei present were distributed more regularly along the fiber axis, reducing the prevalence of cytoplasmic regions disproportionately remote from a myonucleus. This spatial regularization may limit local disparities in access to myonuclear gene products as the cytoplasmic volume associated with each nucleus increases. Importantly, this organization was expressed primarily along the fiber’s long axis. When evaluated across the full fiber surface, the more regular longitudinal arrangement accounted for nearly all of the increase in spatial uniformity, whereas circumferential positioning made only a minimal additional contribution. Thus, the elongated geometry of a myofiber makes longitudinal organization particularly important for limiting nucleus-distant cytoplasmic regions when fewer myonuclei are available relative to fiber size.

## Introduction

Skeletal muscle fibers present an unusual challenge for intracellular organization. A single fiber can extend over millimeters or centimeters while maintaining a continuous cytoplasm that is orders of magnitude larger than that of most mononucleated cells (1-3). These dimensions can separate sites of transcription from the cytoplasmic regions in which the resulting gene products are required over distances rarely encountered in mononucleated cells. The scale of this separation is illustrated by three-dimensional analyses of living mouse fibers, in which mean distances from sampled locations on the sarcolemma or within the cytoplasm to the nearest myonucleus were approximately 20–23 µm, with some regions located more than 50 µm away (4). Multinucleation mitigates this spatial challenge by distributing hundreds of myonuclei throughout the fiber (3), thereby establishing multiple sites of transcriptional output across its large cytoplasmic volume (5-7).

The spatial effectiveness of such a distributed multinucleated system depends on transport distance. Diffusive transport becomes progressively slower as the distance traveled increases, with characteristic diffusion time scaling approximately with the square of distance, whereas the travel time of directed transport scales approximately linearly with distance when mean transport velocity is treated as constant (4, 8). The relative contributions of passive diffusion and active transport therefore depend on the molecular species, transport distance, effective diffusion coefficient, and available intracellular transport machinery. Measurements in skeletal muscle show that protein diffusion is strongly dependent on both the molecule and fiber type (9-11), while microtubule-dependent transport contributes to the intracellular movement of mRNAs and associated translational machinery (12, 13). Passive diffusion and directed transport should therefore be considered complementary processes, both of which remain constrained by the distances between nuclear sources and cytoplasmic targets. Differences in myonuclear number and positioning can substantially alter these distances within the muscle fiber (4, 7, 14, 15).

Spatial proximity may be particularly important because RNAs and proteins do not necessarily become uniformly distributed after synthesis. Transcripts produced by individual myonuclei can remain preferentially localized near their nucleus of origin (16-19). In skeletal muscle, mRNA distribution is influenced by transcript size and active microtubule-based transport, while association with ribosomes can further constrain where translation occurs (12, 13, 20). Proteins generally occupy broader spatial ranges, but their distribution likewise depends on molecular size, binding interactions, nuclear import, and local molecular sinks (9-12, 21, 22). Dystrophin, for example, occupies finite, overlapping sarcolemmal regions associated with individual myonuclei (22). Myonuclear domains are therefore unlikely to represent sharply bounded anatomical compartments (15, 23). Instead, different molecular species occupy overlapping but distinct spatial ranges around distributed myonuclear positions.

Myonuclear number and positioning therefore represent distinct but related determinants of intracellular distance. Over a given longitudinal fiber extent, nuclear number sets the characteristic spacing between myonuclei, whereas their spatial arrangement determines how individual separations vary around that spacing. Myonuclei are non-randomly distributed in normal mammalian muscle (7, 14). Near-uniform nuclear distributions have likewise been described in Drosophila muscle, where organization changes with cellular geometry and depends on distinct microtubule- and LINC-complex-mediated mechanisms (24-27). The biological importance of this organization is supported by studies showing that perturbation of the machinery responsible for positioning or anchoring myonuclei alters muscle structure and function (28-31). Moreover, myonuclear position remains dynamic in mature fibers: following focal physiological damage, resident myonuclei migrate toward the affected region and support local mRNA delivery and sarcomere reconstruction (32). Complementing these biological observations, computational analyses of mammalian muscle fibers have shown that irregular nuclear arrangements create local regions unusually distant from neighboring myonuclei and that the contribution of nuclear arrangement to surface transport distances increases as myonuclear density declines (4). A lower myonuclear number therefore increases characteristic internuclear spacing without, by itself, determining how individual separations are distributed. Whether experimentally restricting postnatal myonuclear addition is accompanied by altered relative spacing within the same muscle and developmental context remains unknown.

Postnatal muscle growth provides an informative setting in which to address this question. During early development, myofibers grow rapidly while muscle stem-cell fusion adds new myonuclei; substantial fiber growth subsequently continues as the rate of myonuclear addition declines (33-35). In the Δ2w model, inducible deletion of Mymk–which encodes the muscle fusogen Myomaker–in muscle stem cells restricts their fusion with growing myofibers. This intervention limits postnatal myonuclear addition while permitting continued fiber growth and maturation, resulting in approximately 50–55% fewer myonuclei in adult fibers (3, 36). By partially uncoupling myonuclear addition from cytoplasmic growth, the model experimentally perturbs the relationship among nuclear number, cell size, and intracellular geometry within the same muscle and developmental context.

The strongly anisotropic geometry of skeletal muscle fibers suggests that spatial adaptation to a smaller myonuclear population may not be expressed equally in all dimensions. The longitudinal dimension greatly exceeds the transverse dimensions, and nuclear movements during muscle development have a pronounced axial component before mature peripheral distributions are established (37-40). Recent in vivo analyses indicate that most myonuclei have reached the fiber periphery by P1 and that nuclei added after P7 remain peripheral rather than first moving through the fiber center (41). Longitudinal positioning may therefore contribute disproportionately to myonuclear organization across the fiber surface. However, whether surface regularity can be explained by longitudinal positioning alone or requires additional circumferential organization has not been tested.

Here, we asked whether restricting postnatal myonuclear addition alters not only the distances between nuclei but also how those distances are distributed as fibers grow. We distinguished the expected increase in absolute internuclear distance associated with fewer nuclei from changes in spacing regularity relative to each fiber’s nuclear number and longitudinal extent. We then tested whether Δ2w fibers exhibited more uniform longitudinal spacing, a lower prevalence of disproportionately large local separations, and pairwise organization centered on the fiber-specific spacing scale. Finally, nested surface null models were used to determine whether genotype differences in organization across the fiber surface could be attributed to longitudinal positioning alone or additionally involved circumferential positioning. The experimental model, analytical framework, and biological hypotheses are summarized in Fig. 1.

**Fig. 1.**
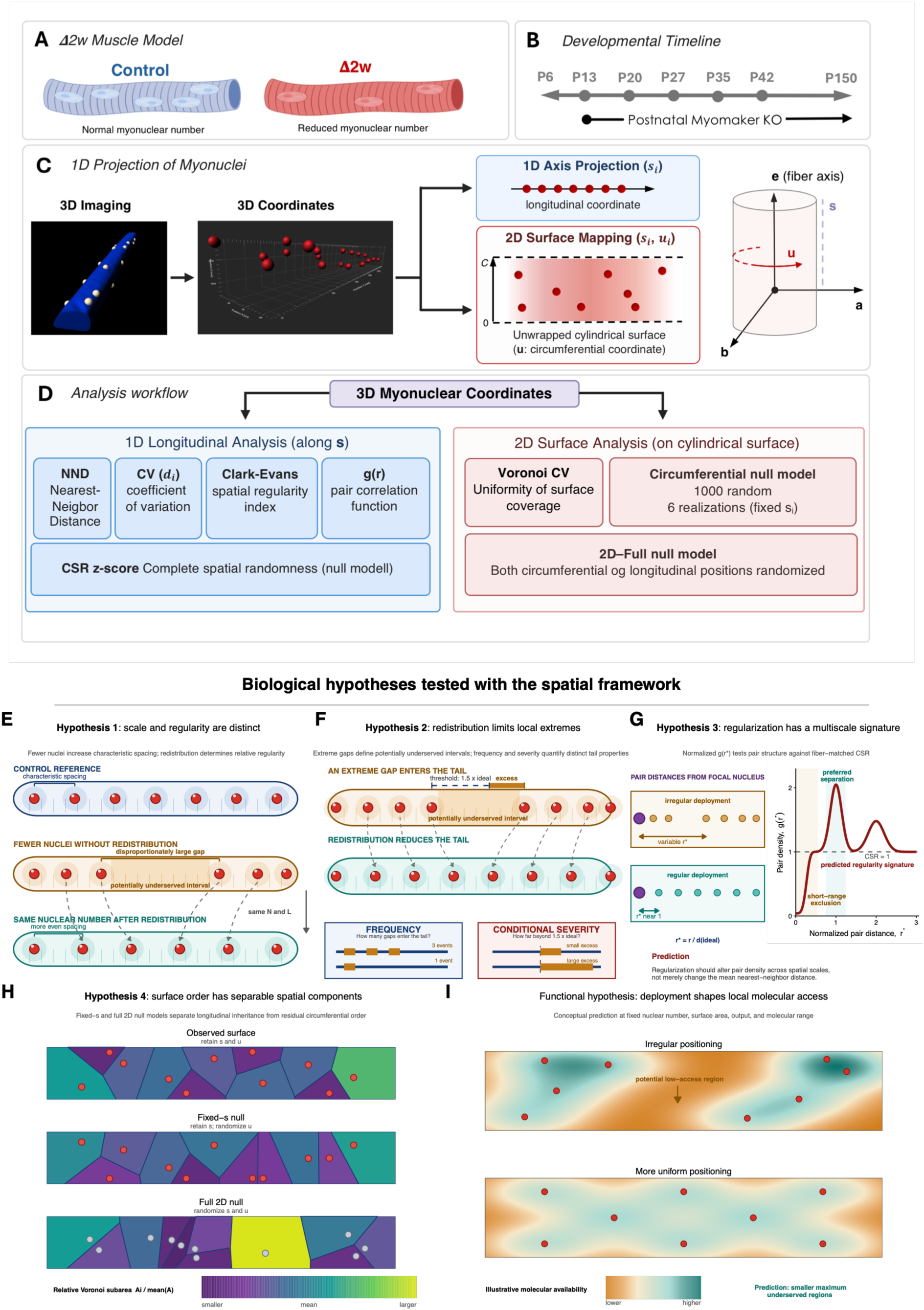
Experimental model and spatial framework for testing myonuclear organization during postnatal growth. **(A)** Schematic of the Δ2w model, in which inducible Myomaker deletion restricts muscle stem-cell fusion and thereby limits postnatal myonuclear addition relative to Control muscle. **(B)** Developmental stages included in the study, spanning postnatal growth from P13 to P150 following postnatal Myomaker deletion. **(C)** Representation of three-dimensional myonuclear coordinates for spatial analysis. Myonuclear positions were analyzed either as longitudinal coordinates, *s_i_*, along the fiber axis or as two-dimensional surface coordinates, (*s_i_*, *u_i_*), after unwrapping the cylindrical fiber surface, where *u_i_* denotes circumferential position. **(D)** Overview of the analytical framework. Longitudinal organization was evaluated using nearest-neighbor distance, the coefficient of variation of adjacent internuclear distances, the Clark–Evans regularity index, and the pair-correlation function *g*(*r*). Fiber-matched complete spatial randomness (CSR) was used to generate null-standardized longitudinal metrics. Surface organization was quantified from the coefficient of variation of Voronoi-partition areas. A fixed-*s* null model preserved longitudinal positions while randomizing circumferential coordinates, whereas the full two-dimensional null model randomized both coordinates. **(E)** Hypothesis 1: reduced myonuclear number increases the characteristic spacing scale but does not uniquely determine relative spatial regularity. The two lower fibers have identical nuclear number and length, and therefore the same fiber-specific spacing scale, yet differ in how individual separations are distributed around that scale. Transparent halos schematically represent local nuclear coverage, and the highlighted interval illustrates a potentially low-access region associated with irregular nuclear positioning. **(F)** Hypothesis 2: redistribution of the remaining nuclei reduces the upper tail of the normalized longitudinal gap distribution. Consecutive gaps were normalized to the fiber-specific ideal spacing as 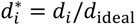, and an extreme gap was defined as 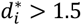. Extreme-gap frequency quantifies the proportion of gaps exceeding this threshold, whereas conditional severity quantifies the mean exceedance beyond the threshold among gaps that enter the tail. **(G)** Hypothesis 3: longitudinal regularization produces a multiscale spatial signature. The normalized pair-correlation function, *g*(*r*^∗^), compares pair density with fiber-matched CSR and can reveal short-range exclusion, enrichment around a characteristic relative spacing, and higher-order spatial organization beyond the nearest-neighbor scale. **(H)** Hypothesis 4: surface regularity can arise either from longitudinal organization alone or from additional circumferential organization. This distinction was tested by comparing the observed Voronoi partition with fixed-*s* and full two-dimensional null configurations. Color denotes Voronoi cell area relative to the mean cell area, 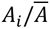. **(I)** Conceptual functional interpretation of myonuclear positioning. At fixed nuclear number, surface area, total molecular output, and characteristic molecular range, irregular positioning is predicted to generate larger regions of relatively low molecular availability, whereas a more uniform spatial distribution is predicted to reduce the maximum extent of a potentially low-access region. Panels E–I illustrate the biological hypotheses and expected spatial signatures and do not represent quantitative experimental results.

## Results

Before examining myonuclear organization, we compared the number of myonuclei per unit fiber length with fiber cross-sectional area across postnatal development. Δ2w fibers contained fewer myonuclei per millimeter than Control fibers at the earliest age examined, and this difference persisted throughout development (Fig. 2A). By contrast, cross-sectional area overlapped substantially between genotypes during early postnatal growth, with lower values in Δ2w fibers becoming clearly apparent only from P35 onward (Fig. 2B). Thus, the difference in longitudinal myonuclear density preceded a clear divergence in fiber caliber, providing a developmental context in which nuclear organization could be examined before and after fiber geometry diverged.

**Fig. 2.**
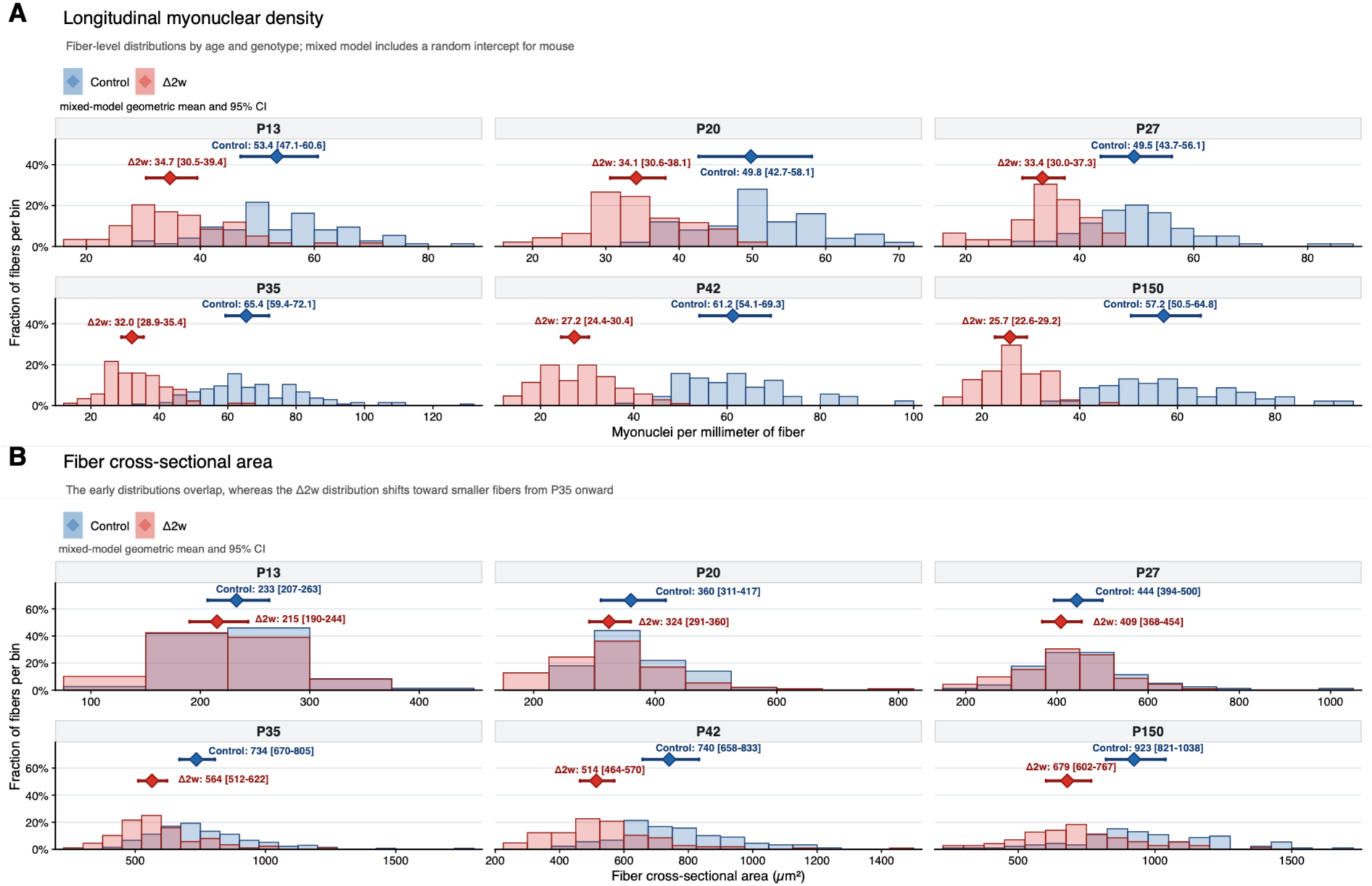
Reduced longitudinal myonuclear density precedes the developmental divergence in fiber cross-sectional area. Fiber-level distributions are shown separately for Control (blue) and Δ2w (red) at each developmental age. Histograms show the fraction of fibers within each bin and were normalized separately within each age-by-genotype group. Diamonds and horizontal whiskers denote mixed-effects-model estimates of the geometric mean and corresponding 95% confidence intervals; numerical estimates and confidence intervals are shown above each distribution. Models were fitted to log-transformed outcomes with age, genotype, and their interaction as fixed effects and a random intercept for mouse to account for clustering of multiple fibers within the same animal. **(A)** Longitudinal myonuclear density, calculated as the number of myonuclei within the analyzed fiber segment divided by segment length and expressed as myonuclei per millimeter. Histogram bin width was 4 myonuclei per mm. **(B)** Fiber cross-sectional area (CSA), calculated as analyzed fiber volume divided by segment length and expressed in µm^2^. Histogram bin width was 75 µm^2^. Axis ranges vary among developmental ages to resolve the within-age distributions. Analyses included 1,029 fibers from 42 mice, with mouse treated as the biological replicate for model-based inference.

These developmental trajectories define the spatial context for the subsequent analyses. Early in development, Δ2w fibers contained fewer myonuclei per unit length despite substantial overlap in fiber caliber. This difference persisted as fiber geometry later diverged, allowing myonuclear organization to be evaluated across distinct combinations of myonuclear number and fiber growth.

### Δ2w fibers exhibit greater longitudinal spacing regularity

For a given longitudinal fiber extent, lower myonuclear density increases the characteristic distance between nuclear positions but does not determine how individual separations vary around that scale. We therefore first asked whether the reduced myonuclear population in Δ2w fibers was accompanied by a change in overall longitudinal regularity. Consecutive-gap variability and nearest-neighbor spacing were evaluated together with fiber-specific complete-spatial-randomness (CSR) references that preserved nuclear number and longitudinal extent (Fig. 3A). These complementary measures distinguish variation in nuclear organization from differences in the spatial scale available to each fiber.

**Fig. 3.**
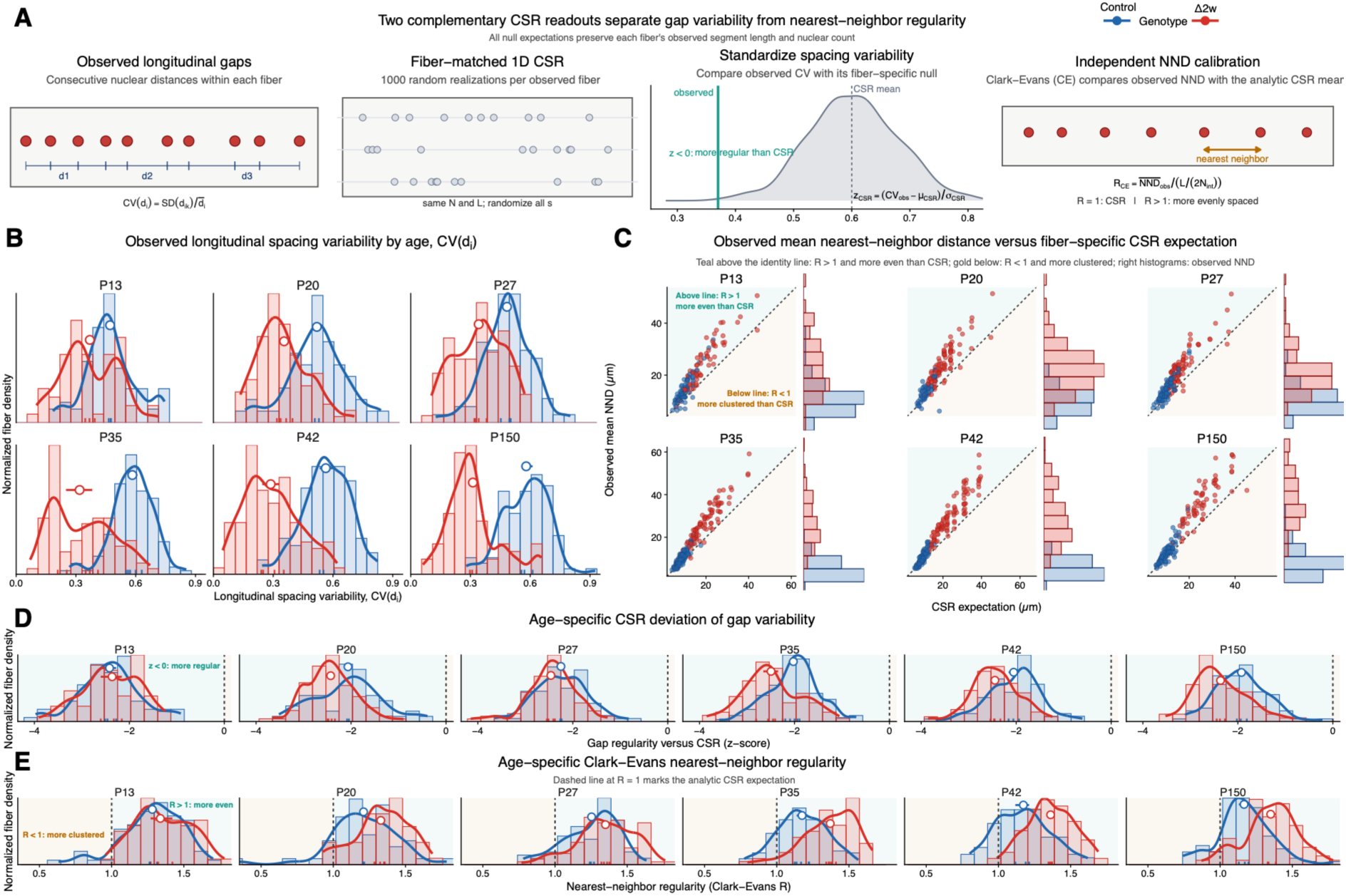
Reduced myonuclear number is accompanied by increased longitudinal spacing regularity. **(A)** Schematic overview of the longitudinal spacing measures and their calibration against one-dimensional complete spatial randomness (CSR). Myonuclei were ordered along the fiber-specific longitudinal axis, and consecutive gaps were defined as *d_ik_* = *s_i_*_,*k*+1_ −*s_ik_*. Within-fiber spacing variability was quantified as 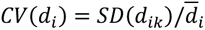, with lower values indicating more uniform consecutive spacing. Fiber-specific CSR distributions were generated by randomizing *N_i_* nuclear positions within the observed segment length *L_i_*. The standardized deviation of observed gap variability from the corresponding CSR distribution was calculated as 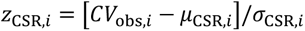; negative values denote lower gap variability than expected for random nuclear positions with the same *N_i_* and *L_i_*. Nearest-neighbor regularity was assessed independently with the one-dimensional Clark–Evans index, 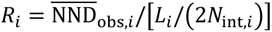, where *N*_int,*i*_ = *N_i_* − 2. *R* = 1 denotes the analytical CSR expectation, *R* > 1 more regular nearest-neighbor spacing, and *R* < 1 relative clustering. **(B)** Age-specific distributions of observed longitudinal gap variability, *CV*(*d_i_*). Histograms and density curves show fiber-level distributions, rug marks denote mouse-level means, and open circles with whiskers show genotype-level means with 95% mouse-bootstrap confidence intervals. **(C)** Observed mean nearest-neighbor distance plotted against the fiber-specific analytical CSR expectation. Each point represents one fiber. The dashed identity line corresponds to *R* = 1; observations above the line have greater mean nearest-neighbor distances than expected under CSR and therefore more uniform nearest-neighbor spacing, whereas observations below the line indicate relative clustering. Marginal histograms show the distributions of observed mean NND. **(D)** Age-specific distributions of the standardized deviation in gap variability from fiber-matched CSR, *z*_CSR_. The dashed line at *z*_CSR_ = 0 denotes agreement with the CSR expectation; increasingly negative values indicate progressively lower spacing variability relative to random nuclear positions with the same nuclear number and segment length. **(E)** Age-specific distributions of the Clark–Evans nearest-neighbor regularity index. The dashed line at *R* = 1 denotes the analytical CSR expectation; values above 1 indicate more regular nearest-neighbor spacing than expected under CSR. Control and Δ2w fibers are shown in blue and red, respectively. Analyses included 968 fibers containing at least six myonuclei and with projected nuclear span no greater than the measured segment length from 42 mice (Control, 482 fibers from 19 mice; Δ2w, 486 fibers from 23 mice). Displayed 95% confidence intervals were obtained from 10,000 mouse-level bootstrap resamples, with the mouse treated as the biological replicate.

The distribution of consecutive internuclear gaps differed markedly between genotypes during postnatal growth (Fig. 3B). Within-fiber gap variability, quantified by *CV*(*d_i_*), was lower in Δ2w fibers than in Control fibers, with the separation becoming particularly pronounced after the earliest developmental stage. Whereas Control fibers showed progressively broader and more variable longitudinal spacing during later growth, Δ2w fibers retained comparatively uniform consecutive spacing. Thus, despite containing fewer nuclei and consequently greater absolute internuclear distances, Δ2w fibers exhibited more uniform longitudinal spacing.

The same pattern was evident when nuclear spacing was evaluated relative to the expectation for random nuclear positions with the same fiber-specific nuclear number and segment length. Observed mean nearest-neighbor distances generally exceeded their corresponding CSR expectations, indicating that myonuclei in both genotypes were distributed more evenly than expected from random placement (Fig. 3C). The displacement above the random expectation was greater in Δ2w fibers during later postnatal development, consistent with enhanced nearest-neighbor regularity in Δ2w fibers.

Normalization of consecutive-gap variability to the complete fiber-specific CSR distribution further separated the observed organization from differences in nuclear density and fiber length (Fig. 3D). Both genotypes exhibited negative *z_CSR_* values, demonstrating lower spacing variability than expected under spatial randomness. At P13, the distributions were comparatively similar, whereas the developmental trajectories subsequently diverged. Δ2w fibers remained strongly displaced toward negative values, while Control fibers shifted toward less negative values during later growth. The reduced gap variability in Δ2w fibers therefore persisted after accounting explicitly for the number of nuclei and longitudinal space available within each fiber.

The Clark–Evans index provided an independent assessment based on nearest-neighbor spacing rather than consecutive-gap variability (Fig. 3E). Values were generally above the random expectation of *R* = 1, confirming non-random longitudinal organization in both genotypes. As development progressed, Δ2w fibers exhibited higher Clark–Evans values than Control fibers, indicating that the remaining nuclei were more evenly separated from their nearest neighbors. The convergence of the gap-based and nearest-neighbor measures shows that the genotype difference was not specific to a single definition of spacing regularity.

Together, these measures show that reduced myonuclear number is associated with greater longitudinal regularity relative to the space available within each fiber. However, measures of overall spacing variability do not identify whether this reorganization specifically limits the largest local deviations from that spacing scale.

### Δ2w fibers contain fewer and less severe relative spacing extremes

Overall longitudinal regularity does not reveal how the largest local separations are distributed. A fiber may show relatively low average spacing variability while still containing isolated gaps that are disproportionately large relative to the spacing available to its nuclear population. We therefore examined the upper tail of the longitudinal gap distribution after normalizing each consecutive separation to the fiber-specific equal-spacing reference (Fig. 4A). This allowed the occurrence and magnitude of relative spacing extremes to be distinguished from the greater absolute spacing associated with reduced myonuclear density.

**Fig. 4.**
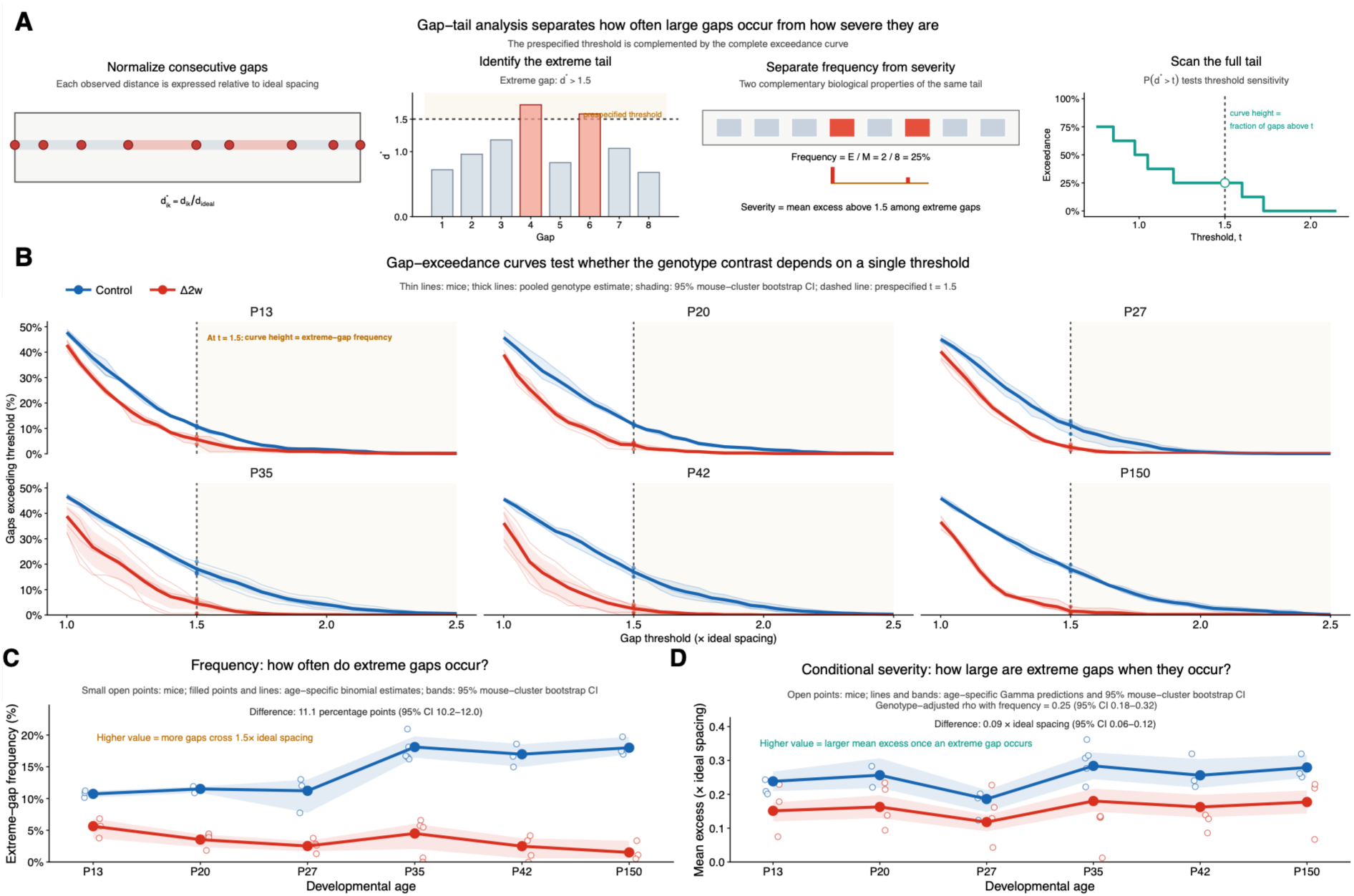
Reduced myonuclear number is associated with fewer and less severe extreme longitudinal gaps. **(A)** Schematic overview of the density-normalized gap-tail analysis. Consecutive longitudinal internuclear gaps, *d_ik_*, were expressed relative to the fiber-specific ideal even-spacing distance, *d*_ideal,*i*_ = *L_i_*/(*N_i_* − 1), yielding 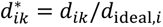 An extreme gap was defined a priori as 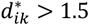. Extreme-gap frequency was defined as the proportion of consecutive gaps exceeding this threshold, whereas conditional severity quantified the mean exceedance above 1.5 among extreme gaps. The exceedance function, *P*(*d*^∗^ > *t*), summarizes the proportion of gaps exceeding each threshold *t* across the upper tail of the distribution. Schematics are illustrative and do not represent individual experimental fibers. **(B)** Age-specific gap-exceedance curves for Control and Δ2w fibers across thresholds from *t* = 1.0 to 2.5. Thin lines show individual-mouse estimates, thick lines show pooled genotype estimates, and shaded regions denote pointwise 95% mouse-cluster bootstrap confidence intervals. The dashed vertical line marks the prespecified threshold *t* = 1.5; the height of each curve at this threshold corresponds to extreme-gap frequency. **(C)** Extreme-gap frequency across postnatal development. Small open circles show mouse-level frequencies. Filled points and connecting lines show age-specific estimates from the grouped binomial model, with shaded regions denoting 95% mouse-cluster bootstrap confidence intervals. The age-standardized difference between genotypes was 11.1 percentage points (95% CI, 10.2-12.0). **(D)** Conditional severity among fibers containing at least one extreme gap. Conditional severity represents the mean exceedance above 1.5 times the fiber-specific ideal spacing. Open circles show mouse-level summaries; lines show age-specific Gamma-model predictions, and shaded regions denote 95% mouse-cluster bootstrap confidence intervals. The age-standardized difference was 0.09 times the ideal spacing (95% CI, 0.06-0.12). Extreme-gap frequency and conditional severity were modestly associated after adjustment for genotype (Spearman’s *ρ* = 0.25; 95% mouse-cluster bootstrap CI, 0.18-0.32). Analyses of extreme-gap frequency included 1029 fibers from 42 mice. Conditional severity was evaluated among the 553 fibers containing at least one extreme gap (448 Control and 105 Δ2w fibers from 40 mice). Control and Δ2w are shown in blue and red, respectively. Confidence intervals were obtained from 10,000 mouse-cluster bootstrap resamples, with mice treated as the biological replicates.

Under this normalization, 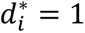 represents the spacing obtained if the available nuclei were distributed evenly across the analyzed longitudinal extent, whereas progressively larger values represent increasing departures from that spacing. Extreme gaps were defined a priori as 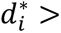 1.5. Their distribution was then characterized by two complementary properties: extreme-gap frequency, which quantified how often gaps crossed this threshold, and conditional severity, which quantified how far beyond the threshold those extreme gaps extended when present (Fig. 4A). The complete exceedance distribution was additionally examined across a range of thresholds to determine whether the genotype difference was specific to the prespecified cutoff.

The upper tail of the normalized spacing distribution differed consistently between genotypes (Fig. 4B). Across postnatal development, a smaller proportion of gaps in Δ2w fibers exceeded a given relative spacing threshold than in Control fibers. This separation extended across a broad range of thresholds rather than emerging only at the prespecified 1.5-times-ideal-spacing cutoff. The genotype difference therefore reflected a broader shift in the upper tail of the relative gap distribution rather than dependence on a single threshold.

At the prespecified threshold, extreme-gap frequency was lower in Δ2w fibers throughout postnatal development (Fig. 4C). The developmental trajectories also diverged with age. Extreme-gap frequency remained comparatively low in Δ2w fibers, whereas Control fibers showed a higher frequency that became more pronounced during later growth. After standardization across developmental age, extreme-gap frequency was 11.1 percentage points lower in Δ2w than in Control fibers (Control - Δ2w,11.1 percentage points; 95% mouse-cluster bootstrap CI,10.2-12.0). Thus, despite their greater absolute internuclear distances, Δ2w fibers were substantially less likely to contain local separations exceeding 1.5 times the spacing defined by their own nuclear number and longitudinal extent.

The magnitude of the extreme gaps that remained was evaluated separately from their frequency (Fig. 4D). Conditional severity quantified how far extreme gaps extended beyond the prespecified threshold and was therefore evaluated only among fibers containing at least one such gap. This analysis included 553 fibers (448 Control and 105 Δ2w) from 40 mice. Across development, extreme gaps in Δ2w fibers showed smaller exceedances than those in Control fibers. The age-standardized difference in conditional severity was 0.09 times the ideal spacing (Control - Δ2w: 0.09, 95% mouse-cluster bootstrap CI, 0.06-0.12). Thus, Δ2w fibers were not only less likely to contain an extreme gap, but the relatively few extreme gaps that occurred also extended less far beyond the fiber-specific spacing threshold.

Extreme-gap frequency and conditional severity captured related but non-equivalent properties of the spacing distribution. After adjustment for genotype, the association between the two measures was modest (Spearman’s *ρ* = 0.25; 95% mouse-cluster bootstrap CI, 0.18-0.32), indicating that fibers in which extreme gaps occurred more frequently tended to show somewhat larger exceedances, but that gap occurrence did not determine their conditional magnitude.

These analyses show that the genotype difference in longitudinal organization extends to the upper tail of the local gap distribution, but consecutive gaps describe only relationships between adjacent nuclei. Whether the same organization is evident across broader pairwise distance scales therefore remained unresolved.

### Myonuclear pairs concentrate near the fiber-specific spacing scale in Δ2w fibers

The preceding analyses showed that Δ2w fibers combined a reduced myonuclear population with greater longitudinal regularity and fewer disproportionately large gaps. These measures, however, primarily describe relationships between adjacent nuclei and do not reveal whether the observed organization extends across broader pairwise distance scales. We therefore examined the one-dimensional pair-correlation function, *g*(*r*), which quantifies the relative frequency of nuclear pairs separated by a given distance. This analysis distinguishes depletion of pairs at short separations from enrichment around characteristic pairwise distances.

Because absolute pairwise distances depend on both nuclear number and longitudinal fiber extent, direct comparison of *g*(*r*) between fibers would confound spatial organization with differences in spacing scale. Pairwise distances were therefore expressed relative to the fiber-specific equal-spacing reference, yielding *g*(*r*^∗^). Under this normalization, *r*^∗^ = 1 corresponds to the separation obtained if the observed nuclei were distributed evenly across the analyzed longitudinal extent, allowing pairwise organization to be compared on a common relative scale.

The spatial signatures resolved by *g*(*r*^∗^) are illustrated schematically in Fig. 5A. A lack of spacing-specific organization produces little enrichment at any particular relative separation, whereas increasing regularity is expected to produce depletion at short distances together with enrichment around *r*^∗^ = 1.

**Fig. 5.**
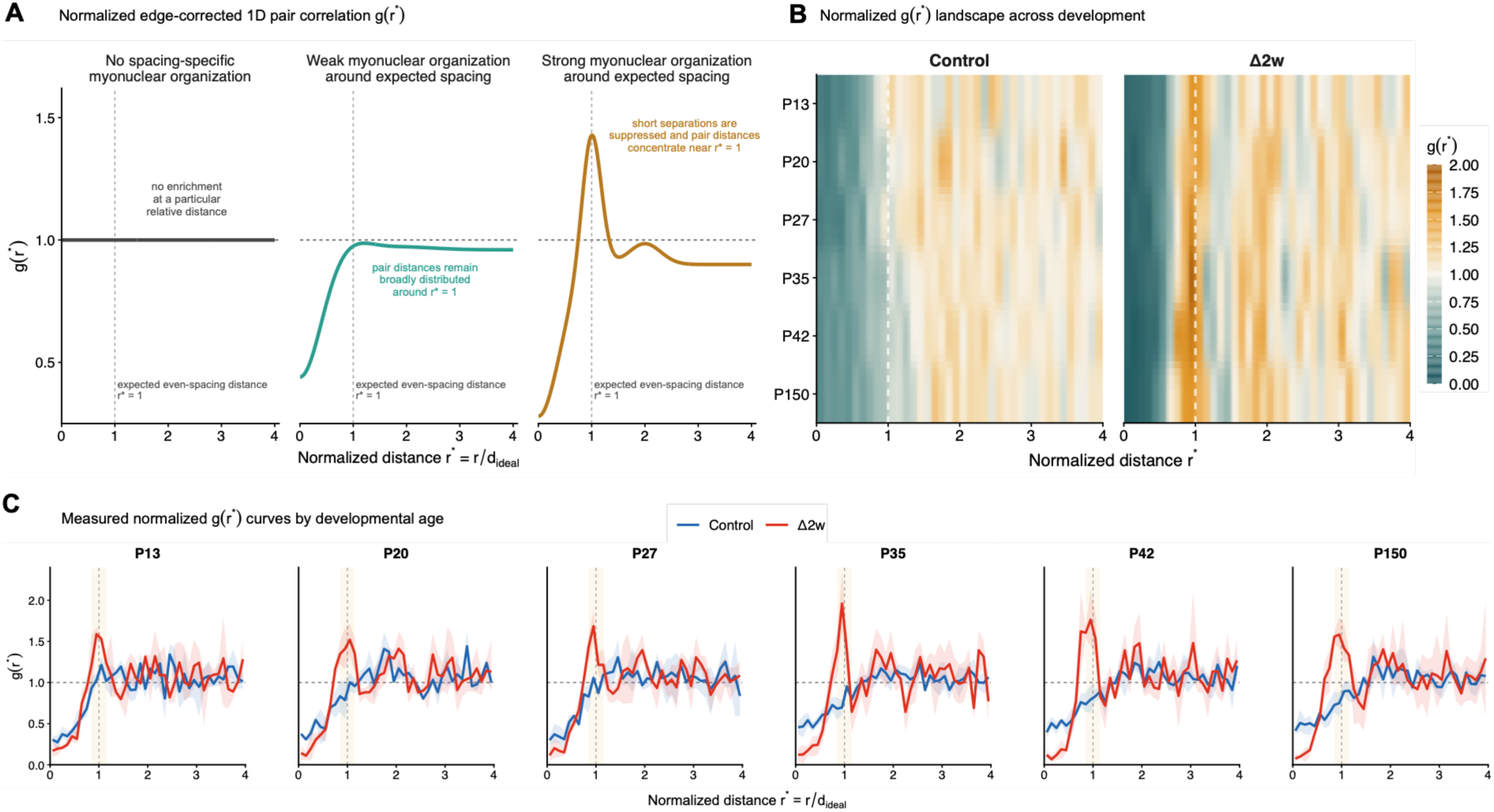
Normalized pair correlation reveals myonuclear organization around a fiber-specific spacing scale. **(A)** Conceptual interpretation of the normalized, edge-corrected one-dimensional pair correlation function *g*(*r*^∗^), where *r*^∗^ = *r*/*d*_ideal_ and *d*_ideal_ = *L*/(*N* − 1). A flat profile at *g*(*r*^∗^) = 1 indicates no enrichment of nuclear pairs at a particular relative separation. Depletion at short distances combined with a broad distribution around *r*^∗^ = 1 indicates weak spacing-specific organization, whereas a pronounced peak near *r*^∗^ = 1 indicates that nuclear-pair distances are concentrated around the expected even-spacing distance. **(B)** Developmental landscapes of normalized *g*(*r*^∗^) for Control and Δ2w fibers. Rows represent postnatal ages from P13 to P150, and color indicates the relative frequency of nuclear pairs at each normalized separation. The vertical dashed line marks *r*^∗^ = 1. **(C)** Age-resolved normalized *g*(*r*^∗^) curves. Solid lines show equal-weight means of mouse-level curves, and shaded bands denote pointwise 95% confidence intervals obtained from 10,000 mouse-cluster bootstrap samples. The vertical dashed line marks *r*^∗^ = 1, and the highlighted interval, 0.85 ≤ *r*^∗^ ≤ 1.15, was used to summarize enrichment around the expected even-spacing distance. Values below and above 1 indicate, respectively, depletion and enrichment relative to conditional fixed-*N*complete spatial randomness within the measured fiber window. The analysis included 982 fibers (482 Control and 500 Δ2w fibers) from 42 mice. Forty-seven fibers in which the projected nuclear span exceeded the measured segment length were excluded.

After normalization to the fiber-specific spacing scale and equal weighting of animals, the pair-correlation landscape differed markedly between Control and Δ2w fibers around *r*^∗^ = 1 (Fig. 5B). Age-resolved curves confirmed this pattern: at every developmental age, Δ2w fibers displayed a localized first peak at *r*^∗^ = 0.95 − 1.05, whereas the corresponding Control profiles were broader and less consistently centered on the reference spacing (Fig. 5C).

To quantify the feature shown in Fig. 5C without performing pointwise tests across the complete curve, we averaged *g*(*r*^∗^) over the prespecified interval 0.85 ≤ *r*^∗^ ≤ 1.15. Pair enrichment within this interval was greater in Δ2w fibers at every age. The mouse-balanced Δ2w–Control difference was 0.377 at P13 (95% mouse-cluster bootstrap CI, 0.315–0.431), increased to 0.689 at P42 (0.573–0.808), and remained 0.664 in adulthood at P150 (0.540–0.768). The age-specific confidence intervals excluded zero at all six developmental stages.

Δ2w fibers additionally showed stronger depletion of myonuclear pairs at short relative separations, followed by sharply localized enrichment around *r*^∗^ = 1 (Fig. 5C). Because the estimator conditions on each fiber’s observed nuclear number and uses the same fixed segment window for distance normalization, edge correction, and CSR calibration, this pattern cannot be attributed solely to the lower nuclear number or to an *N*-dependent shift in the estimator’s baseline. Instead, reduced myonuclear number was associated with a tighter concentration of pairwise separations around the spacing scale defined by each fiber’s own length and nuclear number.

Thus, the Δ2w phenotype reflects not only reduced variability among consecutive gaps, but a broader pairwise organization centered on the fiber-specific spacing scale. The remaining question is whether this one-dimensional organization is sufficient to account for the spatial pattern observed across the two-dimensional fiber surface.

### Longitudinal positioning accounts for nearly all myonuclear surface regularity in Δ2w fibers

Together, the longitudinal analyses showed that reduced myonuclear number was accompanied by greater regularity across several spatial scales, from consecutive gaps to broader pairwise organization. Because myonuclei are positioned predominantly in a subsarcolemmal layer near the fiber surface, we next asked whether the longitudinal organization identified above was sufficient to account for their two-dimensional peripheral distribution, or whether additional organization around the fiber circumference was required. We therefore compared the observed surface organization with two nested null models that separated longitudinally inherited structure from residual circumferential organization (Fig. 6A). The analysis included 968 fibers from 42 mice.

**Fig. 6.**
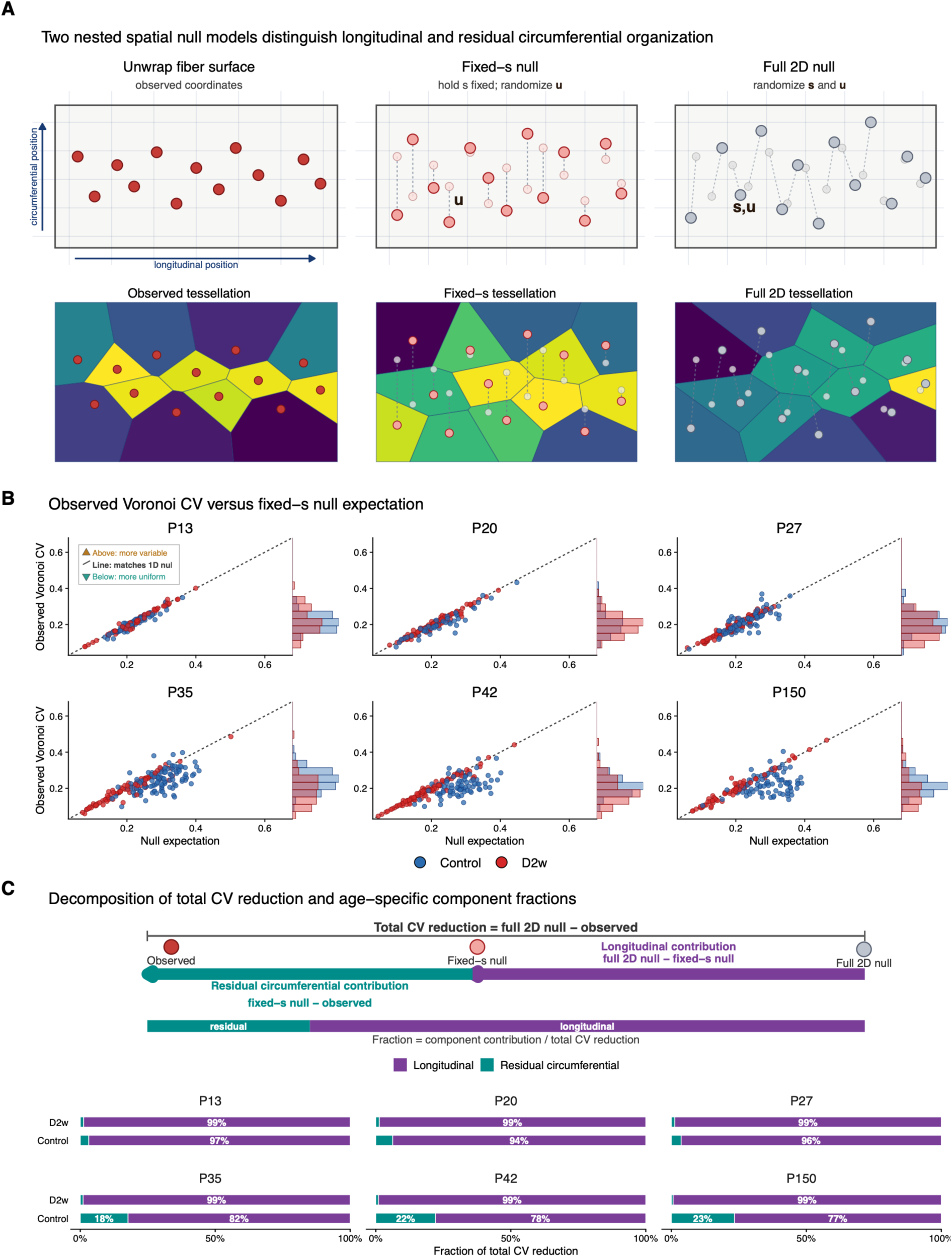
Longitudinal positioning accounts for most surface regularity in Δ2w fibers. **(A)** Schematic representation of the observed myonuclear coordinates on an unwrapped fiber surface and the two nested spatial null models. In the fixed-*s* null model, the observed longitudinal coordinates, *s_i_*, were retained while circumferential coordinates, *u_i_*, were independently randomized. In the full 2D null model, both longitudinal and circumferential coordinates were randomized within the same fiber-specific surface window. The lower row illustrates the corresponding periodic, window-clipped Voronoi partitioning. Schematics are illustrative and do not represent individual experimental fibers. **(B)** Observed Voronoi cell-area coefficient of variation (Voronoi CV) plotted against the mean expectation from the fixed-*s* null model, stratified by developmental age. Each point represents one fiber; Control and Δ2w fibers are shown in blue and red, respectively. The dashed identity line denotes equality between the observed value and the null expectation. Values below the line indicate more uniform Voronoi partitioning than expected from longitudinal organization alone, whereas values above the line indicate greater surface heterogeneity. The interpretation key in the P13 panel applies to all ages. Marginal histograms show the genotype-specific distributions of observed Voronoi CV. **(C)** Additive decomposition of the reduction in Voronoi CV relative to the full 2D null model. Total regularity was defined as Δ_total_ = *μ*_full 2D_ − *CV*_obs_, the longitudinal contribution as Δ_long_ = *μ*_full 2D_ − *μ*_fixed-_*_s_*, and residual circumferential organization as Δ_circ_ = *μ*_fixed-_*_s_* − *CV*_obs_, such that Δ_total_ = Δ_long_ + Δ_circ_. Stacked bars show the longitudinal and residual circumferential fractions of the total reduction for each age and genotype. Fractions were calculated from mouse-balanced group means. Ninety-five percent confidence intervals were obtained from 10,000 mouse-level bootstrap samples but are not displayed in the stacked bars.

Observed surface Voronoi CV closely tracked the expectation generated by the fixed-*s* null model across development (Fig. 6B). This correspondence was particularly pronounced in Δ2w fibers, for which observed values remained close to the identity line across ages. Thus, randomizing circumferential positions while preserving the measured longitudinal arrangement reproduced most of the surface heterogeneity present in the observed Δ2w distributions. In Control fibers, observed Voronoi CV also closely followed the fixed-s expectation at early developmental ages, whereas an increasing proportion of fibers fell below the fixed-*s* expectation at later ages. Values below the identity line indicate more uniform surface partitioning than predicted from longitudinal organization alone and therefore identify an additional contribution from circumferential organization.

To quantify these dimensional contributions, the total reduction in Voronoi CV relative to complete two-dimensional randomness was decomposed additively into a longitudinal component and a residual circumferential component (Fig. 6C). The longitudinal contribution was defined as Δ_long_ = *μ*_full 2D_ − *μ*_fixed-_*_s_* whereas residual circumferential organization was defined as Δ_circ_ = *μ*_fixed-_*_s_* − *CV*_obs_. Together, Δ_total_ = Δ_long_ + Δ_circ_.

In Δ2w fibers, the longitudinal component represented about 99% of the total reduction in Voronoi CV at each developmental age examined. The corresponding longitudinal fractions in Control fibers were 97%, 94%, and 96% at P13, P20, and P27, respectively, but decreased to 82%, 78%, and 77% at P35, P42, and P150. Consequently, residual circumferential organization contributed only approximately 1% of the total CV reduction in Δ2w fibers throughout development, whereas its contribution in Control fibers increased to 18%, 22%, and 23% at P35, P42, and P150, respectively.

The increased surface regularity observed in Δ2w fibers was therefore not accompanied by a corresponding increase in circumferential organization. Instead, the two-dimensional surface pattern in Δ2w fibers was accounted for almost entirely by the observed longitudinal arrangement of the nuclei. Control fibers showed a different developmental pattern, with an additional circumferential contribution becoming evident at later ages. Thus, the spatial organization associated with experimentally restricted postnatal myonuclear addition was predominantly longitudinal rather than reflecting a generalized increase in regularity across both surface dimensions.

## Discussion

The central finding of this study is not simply that fewer myonuclei are separated by greater distances; that consequence follows directly from nuclear number and fiber size. Rather, experimentally restricted postnatal myonuclear addition altered how those greater distances were distributed during fiber growth. After calibration against random patterns preserving each fiber’s nuclear number and longitudinal extent, Δ2w fibers showed greater longitudinal spacing regularity than Control fibers (Fig. 3B–E; Fig. 7B). They also contained fewer and less severe separations that were disproportionately large relative to their own spacing scale (Fig. 4B–D; Fig. 7C). Biologically, these analyses distinguish a nuclear population that is merely farther apart from one that is also distributed more evenly within the longitudinal space available to the fiber. Myonuclear number constrains the characteristic scale of nuclear separation, whereas nuclear arrangement determines whether those separations are distributed relatively uniformly or concentrated into clusters and nucleus-distant regions (Fig. 7A) (4, 7, 14, 15). In biological terms, a sparse but regular nuclear population may limit local disparities in access to myonuclear gene products more effectively than an equally sparse but irregular population (Fig. 7E).

**Fig. 7.**
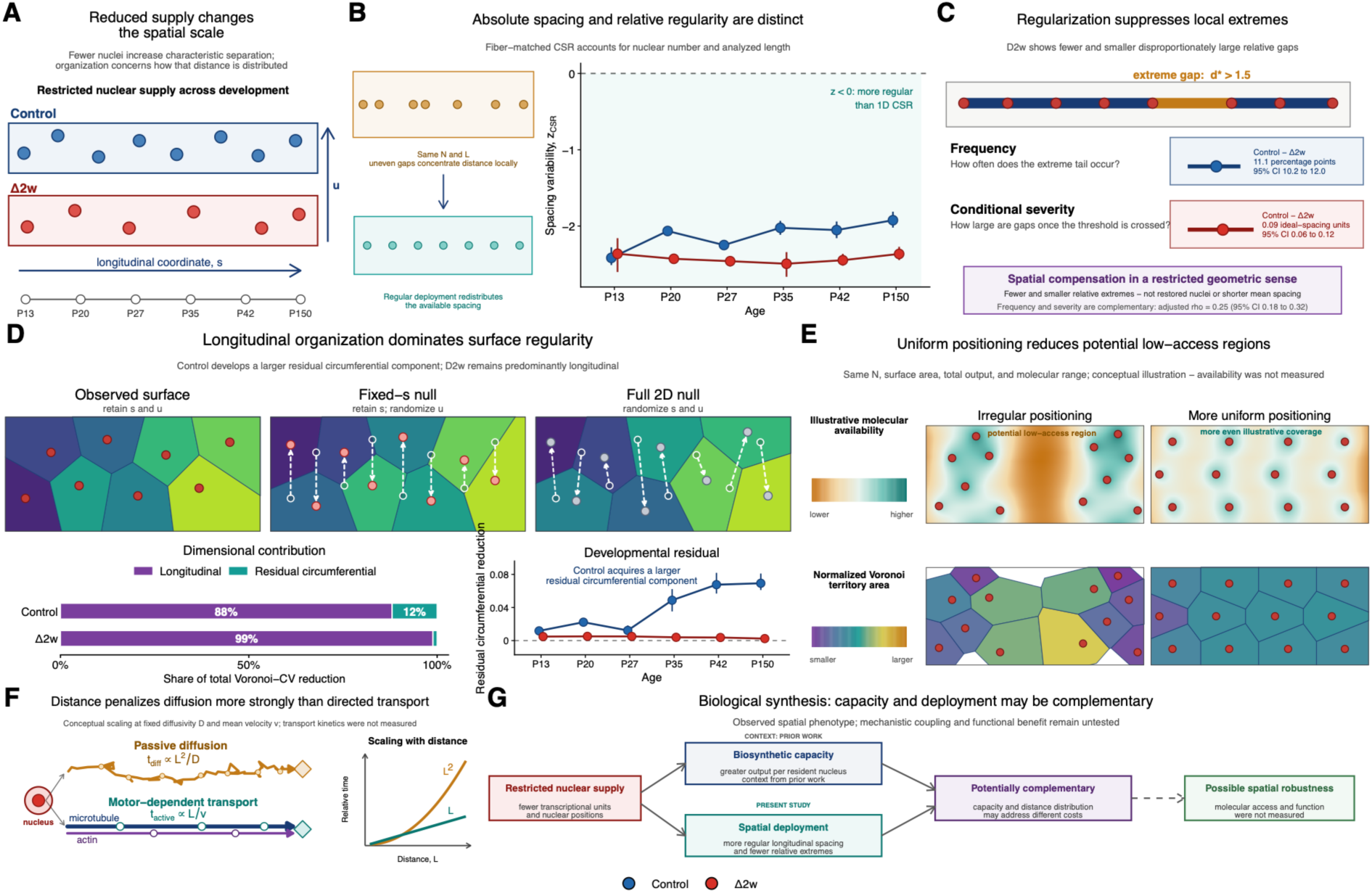
Spatial organization of a reduced myonuclear population and its potential biological implications. **(A)** Conceptual distinction between the spatial scale imposed by myonuclear number and the organization of nuclei within that scale. Reducing myonuclear number increases the characteristic separation between the remaining nuclear positions. The schematics illustrate this consequence along the longitudinal coordinate, *s*, while indicating the circumferential coordinate, *u*, used in the surface analyses. Redistribution of the remaining nuclei can alter the relative uniformity of their spacing without replacing myonuclei that were not added during growth or reducing the characteristic separation imposed by nuclear number. **(B)** Longitudinal spacing regularity after accounting for fiber-specific nuclear number (*N*) and analyzed fiber segment length (*L*). The schematic illustrates how fibers with identical *N* and *L* can differ in spatial organization when the available distance is concentrated into irregular gaps or distributed more evenly. Observed consecutive-gap variability was standardized relative to a fiber-matched one-dimensional complete spatial randomness (CSR) reference as *z*_CSR_. Negative values indicate lower spacing variability, and therefore greater longitudinal regularity, than expected for random nuclear positions with the same nuclear number and longitudinal extent. Points show age-specific genotype estimates and error bars denote 95% confidence intervals obtained by mouse-level bootstrap resampling. **(C)** Relative spacing extremes under reduced myonuclear number. Normalizing consecutive gaps to the fiber-specific ideal spacing distinguishes expected increases in spacing from disproportionately large local separations. Δ2w fibers showed fewer and less pronounced extreme gaps, indicating that the increased spacing associated with reduced nuclear number was distributed with fewer local spatial extremes. **(D)** Dimensional origin of myonuclear surface regularity. Observed surface coordinates retain both longitudinal and circumferential nuclear positions. In the fixed-*s* null model, measured longitudinal positions are retained while circumferential positions are randomized; in the full 2D null model, both coordinates are randomized within the same fiber-specific surface geometry. Open white circles indicate original positions, filled circles randomized positions, and dashed arrows the corresponding displacements. Voronoi colors distinguish individual Voronoi cells and do not encode Voronoi cell area. The reduction in Voronoi cell-area CV relative to the full 2D null was partitioned into longitudinal and residual circumferential components. After standardization across developmental ages, longitudinal organization accounted for 88% of the total reduction in Control fibers and 99% in Δ2w fibers, leaving residual circumferential contributions of 12% and approximately 1%, respectively. Age-resolved estimates show that the residual circumferential contribution increased during later development in Control fibers but remained small in Δ2w. **(E)** Conceptual illustration of how different myonuclear patterns could influence spatial access to nucleus-derived molecular products. The two configurations contain the same number of nuclei and assume identical surface area, total molecular output, and characteristic molecular range. The upper maps depict an illustrative distance-dependent availability field, with orange indicating lower and teal higher relative availability. Irregular positioning produces greater spatial heterogeneity and a potential low-access region, whereas more uniform positioning distributes the illustrative field more evenly. The lower maps show the corresponding Voronoi partitioning, with Voronoi cell area normalized to the mean Voronoi region, *A_i_*/(*A*_surface_/*N*); purple denotes smaller and gold larger relative Voronoi cell areas. These panels are conceptual and do not represent measured transcript abundance, protein distribution, molecular transport, or local functional output. **(F)** Conceptual comparison of the distance dependence of passive diffusion and directed intracellular transport. For fixed diffusivity, *D*, characteristic diffusion time scales with the square of transport distance, *t*_diff_ α *L*^2^/*D*, whereas transport at a constant mean velocity, *v*, scales approximately linearly with distance, *t*_active_ α *L*/*v*. The schematic therefore illustrates that increasing source-to-target distance imposes a progressively greater time penalty on diffusion than on directed transport. The trajectories and scaling curves are conceptual; molecular diffusivities, motor velocities, and transport kinetics were not measured in the present study. **(G)** Integrated biological model. Restricted satellite-cell fusion reduces the addition of new myonuclei during growth, resulting in fewer myonuclei and a lower spatial density of nuclear positions within the fiber. Previous work indicates that resident myonuclei can increase biosynthetic output when myonuclear number is reduced, whereas the present study identifies a more regular longitudinal distribution and fewer relative spacing extremes. The diagram summarizes biosynthetic capacity and spatial organization as potentially complementary features of fibers containing fewer myonuclei while distinguishing prior experimental context from findings of the present study. Their mechanistic relationship and consequences for local molecular availability or muscle function were not tested. Panels A, E, F, and G are conceptual representations; panels B–D summarize quantitative findings from the present study. Confidence intervals in panels B–D were obtained from 10,000 mouse-level or mouse-cluster bootstrap replicates, as appropriate, with the mouse treated as the biological replicate.

Why might variation in nucleus-to-cytoplasm distance matter? The answer depends on the spatial range of nucleus-derived products. Some muscle transcripts remain enriched near their nucleus of origin, whereas others are actively transported along microtubules and disperse over several tens of micrometres (12, 13, 16-19, 22). Proteins likewise differ in their capacity to disperse through the syncytium as a function of molecular size, localization signals, binding interactions, and turnover (9-11, 22). A large source-to-target distance will therefore not affect every molecular product in the same way. Stable and mobile proteins may cross several nominal myonuclear domains, whereas short-lived, locally captured, or poorly mobile products may remain sensitive to the position of their nucleus of origin.

Dystrophin illustrates this principle particularly clearly. Dystrophin produced by individual myonuclei occupies spatially restricted sarcolemmal regions, and restoration of a sufficient number of overlapping dystrophin domains influences treatment efficacy in dystrophic muscle (22). The cytoplasmic extent over which a myonucleus contributes a particular product is therefore product specific rather than a fixed anatomical compartment. There is no single critical distance beyond which the availability of all nucleus-derived products necessarily becomes insufficient. Instead, each transcript or protein has an effective molecular range determined by its production, transport, stability, and localization. Variation in nuclear spacing becomes biologically relevant when the distance between neighboring nuclear sources approaches or exceeds that range.

The upper tail of the spacing distribution may be particularly relevant biologically. Whereas mean spacing and overall spacing variability describe the typical organization of the nuclear population, the upper-tail analysis identifies individual separations that are disproportionately large relative to the spacing defined by the same fiber’s nuclear number and longitudinal extent (Fig. 4B–D; Fig. 7C). These separations delineate intervening fiber regions that are unusually remote from neighboring myonuclei and may therefore be most sensitive to limitations in the intracellular range of locally required transcripts or proteins. Previous three-dimensional analyses of living EDL and soleus fibers found mean nearest-nucleus distances of approximately 20–23μm but also identified small regions located more than 50μm from the nearest myonucleus (4). Such regions make a limited contribution to the mean while potentially imposing a substantially greater local transport requirement.

For diffusion, the characteristic time required to cover a distance increases approximately with the square of that distance, whereas active transport at a constant mean velocity scales approximately linearly (Fig. 7F) (4, 8, 15, 42, 43). Under otherwise fixed conditions a twofold increase in distance therefore produces a fourfold increase in the temporal cost of diffusion, which may influence how myofibers maintain size homeostasis on a temporal scale (44). These considerations do not establish that the largest myonuclear separations produce local molecular deficits. They do, however, explain why the upper tail deserves separate consideration. If molecular availability declines with distance from a producing nucleus, concentrating a given total distance into a few large separations will create greater local variation than distributing the same distance more evenly. The lower occurrence and severity of relative spacing extremes in Δ2w fibers may therefore be biologically more consequential than would be inferred from mean internuclear distance alone (Fig. 4B–D; Fig. 7C).

Earlier comparisons between EDL and soleus provide a biological context for this result. In living adult mouse muscle, EDL fibers contain fewer myonuclei per unit length and larger myonuclear domains than soleus fibers, yet their nuclei are positioned more regularly (4, 7, 14). These analyses showed that similar mean distances can coexist with local distance extremes: small regions disproportionately remote from any myonucleus and therefore potentially less accessible to transcripts or proteins with a limited intracellular range. Computational redistribution further indicated that regular nuclear positioning reduced nucleus-to-surface distances more effectively in the sparsely nucleated EDL than in the more densely nucleated soleus, where the greater number of nuclear positions kept most regions relatively close to a nucleus even after randomization (4).

EDL and soleus also differ in oxidative capacity (45-47), contractile phenotype (48, 49), cytoskeletal organization (14), and intracellular diffusion (9), making it difficult to attribute their nuclear patterns to nuclear density alone. The Δ2w model provides a complementary comparison within a single muscle type and strengthens the broader inference that nuclear arrangement becomes increasingly consequential as nuclear density declines.

The upper-tail analysis describes local relationships between consecutive myonuclei but does not establish whether the same organization extends across broader pairwise distance scales. The normalized pair-correlation analysis addressed this question by testing whether nuclear pairs were concentrated around particular distances after all pairwise separations had been expressed relative to the even-spacing distance of the corresponding fiber (Fig. 5B, C). In Δ2w fibers, pairwise separations were more strongly concentrated near the fiber-specific reference distance, with stronger depletion at shorter relative separations. This argues against a simple positioning system that places neighboring nuclei around an invariant separation in absolute units. Instead, the spatial organization appears to scale with the number of nuclei and the longitudinal space within each fiber.

A scale-dependent pattern of this kind is compatible with self-organizing mechanisms in which nuclear position emerges from interactions among nuclei, cytoskeletal forces and cellular boundaries. Studies of developing muscle have identified cellular machinery capable of moving, separating, and anchoring myonuclei. Microtubules, kinesin and dynein motors, and microtubule-associated proteins contribute to nuclear movement and separation (28-30, 37-40). Coupling of nuclei to the cytoskeleton through LINC-, KASH-, nesprin-, and related protein complexes contributes to nuclear movement, spacing, and anchoring (26, 27, 50-52). Perturbing these systems produces distinct positioning defects and can impair muscle structure or function (27, 28, 31, 53). In Drosophila muscle, near-uniform nuclear spreading can emerge through distance-dependent interactions between nuclei and between nuclei and the cellular boundary (24, 25, 54, 55). Such a force-balance system need not encode a preferred absolute separation. Altering either cellular dimensions or nuclear number changes the balance of interactions and therefore the nuclear configuration that emerges.

The present data are compatible with this class of mechanism but do not establish that it operates in Δ2w fibers. Restricting fusion may change the mechanical relationships among resident nuclei, but it may also alter where nuclei are introduced during growth. Newly fused and previously resident nuclei are not necessarily biologically equivalent; they can exhibit distinct transcriptional programs and influence one another during adaptation (5, 6). Moreover, nuclear movement and nuclear addition are both developmentally regulated, and nuclei introduced at different stages may encounter different cytoskeletal and anatomical environments (30, 32-34, 41). Restricted fusion could consequently influence nuclear organization through several routes, including fewer local fusion events, altered cytoskeletal remodeling, or changes in the transcriptional (56) and mechanical state (57) of resident nuclei. Distinguishing these alternatives will require lineage-specific labeling (5) and direct tracking of nuclear positions during postnatal growth.

The dimensional decomposition, however, provides a particularly informative constraint on possible mechanisms. The longitudinal analyses establish how myonuclei are arranged along the fiber axis, but myonuclei occupy a two-dimensional subsarcolemmal surface. The nested surface models therefore addressed a separate biological question: whether the axial arrangement was sufficient to account for the distribution of nuclear positions across the complete fiber surface, or whether additional circumferential organization was required (Fig. 6A–C; Fig. 7D). Nearly all of the surface regularity in Δ2w fibers was retained when the measured longitudinal coordinates were preserved and the circumferential coordinates were randomized. The more even relationship between myonuclei and the fiber surface can therefore be attributed primarily to axial order rather than to a generalized increase in organization across both surface dimensions.

The predominance of axial organization is plausible given the architecture of the fiber and the orientation of its cytoskeleton. Longitudinal microtubule bundles provide tracks for nuclear movement, and nuclear translocation during myotube formation, maturation, and repair has a pronounced component along the fiber axis (30, 40, 58). A change in axial position also alters nuclear relationships over a dimension that may extend for millimetres or centimetres (59), whereas the circumferential coordinate is periodic and spans a much shorter distance. Longitudinal order can consequently produce a more even distribution of nearest-nucleus distances across the fiber surface without requiring a corresponding increase in circumferential order.

Circumferential position should nevertheless not be regarded as biologically irrelevant. Myonuclei interact with blood vessels, desmin networks, and other local structures that may influence their peripheral position (31, 41). Specialized myonuclei at neuromuscular and myotendinous junctions are also positioned according to local functional requirements rather than global spacing uniformity (16, 31, 53, 60, 61). Nuclear organization in muscle is therefore not governed by a universal tendency toward maximal uniformity. Global regularity coexists with functionally specified local clustering and transcriptional specialization. The present decomposition shows specifically that circumferential coordinates made only a minimal additional contribution to the global surface regularity of Δ2w fibers; it does not exclude important circumferential relationships at specialized sites (Fig. 6C; Fig. 7D).

The developmental behavior of Control fibers may also be informative and warrants further study. Longitudinal organization accounted for most surface regularity early in development, whereas a larger circumferential contribution emerged at later ages (Fig. 6C; Fig. 7D). This pattern could reflect progressive maturation of the peripheral cytoskeleton, continued incorporation of myonuclei, or increasing influence of local anatomical cues during postnatal development. However, the corresponding circumferential contribution remained small in Δ2w fibers. This may indicate that circumferential organization depends partly on nuclear density or continued myonuclear addition. Alternatively, when the longitudinal pattern is already comparatively regular, circumferential rearrangement may produce little further reduction in surface heterogeneity. These interpretations remain speculative, but they suggest that longitudinal and circumferential positioning need not be regulated identically or mature on the same developmental schedule.

The spatial findings can also be considered alongside the substantial biosynthetic flexibility previously observed in Δ2w fibers. Despite markedly reduced nuclear content, these fibers can expand the cytoplasmic volume associated with resident myonuclei and preserve considerable baseline function (36). Emerging experimental evidence provides a potential transcriptional basis for this flexibility as increasing Polr2a expression in resident myonuclei elevates mRNA abundance, protein accumulation, and fiber size without a corresponding increase in myonuclear number. Increased RNAPII-dependent transcription also reduced myonuclear accretion during postnatal development and overload-induced growth (62).

These observations are consistent with a transcription-driven model in which resident myonuclei initially accommodate increasing cytoplasmic demand by drawing on transcriptional reserve, whereas additional nuclei become increasingly favorable when transcriptional output can no longer scale sufficiently with fiber volume (63). Within this framework, myonuclear accretion is not dictated by fiber size alone but emerges from the balance between cytoplasmic demand and the transcriptional capacity of the resident nuclear population (3, 36, 62, 63).

Transcriptional flexibility, however, addresses only one consequence of reduced myonuclear number. Increasing the biosynthetic output of each nucleus may compensate partly for having fewer nuclear genomes, but it does not eliminate the geometric separation between those genomes and the cytoplasmic regions in which their products are required. Conversely, a more regular nuclear arrangement can reduce variation in nucleus-to-cytoplasm distance but cannot restore the transcriptional capacity that would have been added through satellite-cell fusion. Biosynthetic capacity and nuclear organization should therefore be regarded as complementary rather than interchangeable properties. The first determines how much molecular output the myonuclear population can generate; the second influences the distances over which that output must be transported or remain locally available (Fig. 7G).

In conclusion, the biological consequences of restricted postnatal myonuclear addition cannot be described by myonuclear number alone. Δ2w fibers contained fewer myonuclei and consequently developed enlarged cytoplasmic domains and greater characteristic internuclear distances. Within that expanded spatial scale, however, the remaining nuclei were arranged more regularly along the longitudinal axis, reducing the prevalence of local regions disproportionately remote from a myonucleus. This longitudinal organization accounted for nearly all of the associated increase in uniformity across the fiber surface, with minimal additional contribution from circumferential positioning. Myonuclear number therefore establishes the characteristic scale of nuclear separation, whereas myonuclear arrangement determines how evenly those separations are distributed. Whether this more even distribution of nucleus-to-cytoplasm distances preserves local molecular availability remains to be established, but it identifies a previously underappreciated dimension of myonuclear-domain plasticity.

## Materials and Methods

### Animals and experimental material

Developing extensor digitorum longus (EDL) myofibers were obtained from mice at postnatal day (P)13, P20, P27, P35, P42, and P150 from previously reported experimental cohorts (3, 36). Three-dimensional myonuclear coordinates and corresponding fiber-level measurements of segment length, volume, and surface area were used in the present study to examine the spatial organization of myonuclei across postnatal development. EDL fibers at these ages are composed predominantly of fast type 2x and 2b fibers (14, 34, 45, 46).

Myomaker was conditionally deleted in muscle satellite cells to limit their contribution of new myonuclei during postnatal growth, as described previously (36). Myomaker^loxP/loxP^ mice were crossed with Pax7CreER mice to permit tamoxifen-inducible recombination in satellite cells. Experimental Myomaker^loxP/loxP; Pax7CreER^ mice are denoted Δ2w, whereas littermate Myomaker^loxP/loxP^ mice served as Controls. All animals received 200 µg tamoxifen by intraperitoneal injection at P6. Both female and male mice were represented. All animal procedures associated with generation of the original material were approved by the Institutional Animal Care and Use Committee at Cincinnati Children’s Hospital Medical Center (36).

EDL muscles were fixed in 2% paraformaldehyde before isolation of individual fibers. Fibers were mounted on Superfrost Plus slides (Thermo Fisher Scientific), coverslipped with DAPI Fluoromount-G (SouthernBiotech) to visualize myonuclei, dried overnight, sealed, and imaged within 48 h.

### Confocal imaging and three-dimensional reconstruction

Image stacks were acquired with an Andor Dragonfly spinning-disk confocal microscope equipped with a 40× oil-immersion objective (numerical aperture 1.3) and a Zyla 4.2 sCMOS camera. The x-y sampling interval was 0.3 × 0.3 µm and the z-step was 1 µm. Excitation wavelengths were 405 nm, 488 nm, and 561 nm, and images were acquired with 2 × 2 pixel binning. Approximately 25–40 fibers were analyzed per muscle, as described for the original dataset (3).

Three-dimensional fiber reconstructions were generated in Imaris 8.3.1 (Bitplane). For each analyzed fiber segment, the exported data contained the x, y, and z coordinates of every identified myonucleus together with nuclear number, segment length, fiber volume, and surface area. These measurements were retained at the individual-fiber level and linked to mouse, slide, and fiber identifiers throughout the analysis.

### Dataset assembly and analysis populations

Raw Imaris exports were imported and merged using unique mouse and fiber identifiers. Genotype and developmental age were assigned from the experimental identifiers, and data-integrity checks verified the one-to-one correspondence between each set of nuclear coordinates and its associated fiber-geometry measurements before spatial analysis.

The full dataset comprised 1,029 fibers (519 Control and 510 Δ2w) from 42 mice and contained 14,946 myonuclei. Longitudinal regularity analyses were restricted to fibers containing at least six nuclei (1,015 fibers) to provide stable fiber-level estimates and to match the minimum nuclear number used for surface analyses. The periodic surface Voronoi analysis imposed the additional geometry criteria described below and included 968 fibers from the same 42 mice. Multiple fibers obtained from one mouse were treated as repeated observations from the same biological replicate.

### Longitudinal coordinate system and absolute spacing

For each fiber, principal component analysis (PCA) was applied to the centered three-dimensional myonuclear coordinates. The first principal component defined the longitudinal unit vector, *e_i_*. The longitudinal coordinate of nucleus k in fiber i was

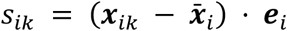

The projected coordinates were ordered from one end of the analyzed segment to the other, and consecutive longitudinal gaps were calculated as

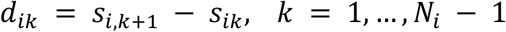

where *N_i_* is the number of myonuclei in fiber i and *L_i_* is the measured Imaris segment length. Nuclear linear density was defined as *N_i_*/*L_i_*. Absolute spacing was summarized by the mean consecutive gap and treated as descriptive geometric context rather than as a measure of relative spatial organization.

### Longitudinal spacing regularity and complete spatial randomness

Within-fiber spacing variability was quantified by the coefficient of variation of consecutive longitudinal gaps,

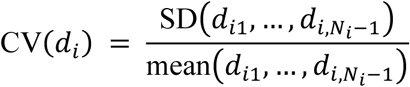

where the sample standard deviation was calculated with *N_i_* − 2 degrees of freedom. Lower CV(*d_i_*) indicates more uniform consecutive spacing within a fiber.

To determine whether the observed spacing variability differed from that expected solely from nuclear number and segment length, each fiber was compared with a fiber-specific one-dimensional complete-spatial-randomness (CSR) reference. For each of 1000 realizations, *N_i_* positions were sampled independently from a uniform distribution on [0, *L_i_*], ordered, and converted to *N_i_* − 1 consecutive gaps. CV (*d_i_*) was calculated identically for each realization. The mean and standard deviation of the simulated distribution, *μ*_CSR,i_ and *σ*_CSR,i_, were used to standardize the observed value:

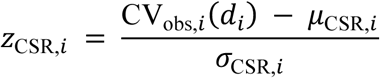

Negative z-scores indicate lower spacing variability, and therefore greater longitudinal regularity, than expected for random points with the same nuclear number and analyzed length.

Longitudinal regularity was also quantified independently with a one-dimensional adaptation of the Clark–Evans index (64). To reduce terminal edge effects, the two outermost nuclei were excluded. For each remaining nucleus, nearest-neighbor distance was the smaller of the distances to its immediate left and right neighbors. If *N*_int,i_ denotes the number of retained interior nuclei, the one-dimensional CSR expectation for the mean nearest-neighbor distance is *L_i_*/(2*N*_int,i_), and the regularity index was

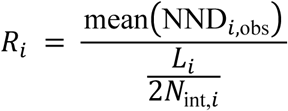

Values of *R_i_* > 1 indicate more regular spacing than expected under CSR, *R_i_* = 1 is consistent with the CSR expectation, and *R_i_* < 1 indicates clustering.

### Density-normalized gap-tail analysis

To characterize the upper tail of the longitudinal gap distribution independently of absolute nuclear density, each consecutive gap was normalized to the fiber-specific even-spacing distance,

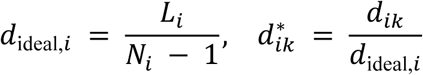

Thus, *d*\**_ik_* = 1 corresponds to the spacing obtained if the observed nuclei were distributed evenly across the analyzed fiber length. An extreme gap was defined a priori as *d*\**_ik_* > 1.5. The threshold was specified before inspection of the exceedance analyses and was not selected from the observed upper-tail curves. For each fiber, the total number of gaps was *M_i_* = *N_i_* − 1 and the number of extreme gaps was

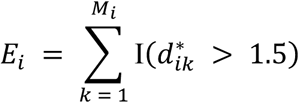

where I(·) is the indicator function. Extreme-gap frequency was *F_i_* = *E_i_*/*M_i_*. Reconstruction from the nuclear coordinates confirmed *M_i_* = *N_i_* − 1 for all 1,029 fibers and reproduced the previously processed fiber-level frequencies to numerical precision.

### Age-resolved extreme-gap probability

Developmental changes in extreme-gap probability were estimated with a grouped binomial generalized linear model with a logit link,

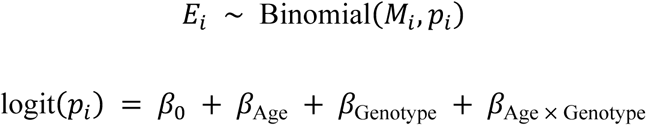

Developmental age was treated as a six-level categorical factor (P13, P20, P27, P35, P42, and P150). The age-by-genotype interaction allowed the genotype difference in extreme-gap probability to vary across development. Model-based probabilities were obtained for each age-by-genotype combination.

Because gaps and fibers sampled from the same animal are not independent biological replicates, uncertainty was estimated by mouse-cluster bootstrap. Mice were sampled with replacement within each age-by-genotype stratum, retaining all fibers and gaps belonging to each selected mouse. Extreme-gap and total-gap counts were then pooled within each bootstrap sample. The procedure was repeated 10,000 times, and pointwise 95% confidence intervals were defined by the 2.5th and 97.5th percentiles of the bootstrap distribution.

### Gap-exceedance curves

Sensitivity to the prespecified threshold was assessed across the complete upper tail of the normalized gap distribution. For each age-by-genotype group, the exceedance probability was calculated as

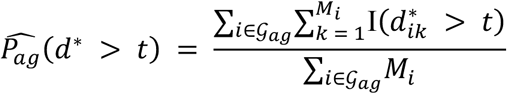

where G*_a_*_g_ denotes the set of fibers belonging to developmental age *a* and genotype *g*, *M_i_* = *N_i_* − 1 is the number of consecutive gaps in fiber *i*, and *I*(⋅) is the indicator function. Thus, 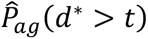 represents the proportion of all consecutive gaps within a given age-by-genotype group that exceed the normalized spacing threshold *t*. Exceedance probabilities were evaluated for thresholds from *t* = 1.0 to 2.5 in increments of 0.05. Pointwise 95% confidence intervals were obtained using the same 10,000-replicate mouse-cluster bootstrap described above. The exceedance curves were used as a threshold-sensitivity analysis; separate hypothesis tests were not performed at each value of *t*.

### Extreme-gap frequency and conditional severity

Extreme-gap behavior was separated into two components: the frequency with which extreme gaps occurred and their magnitude conditional on occurrence. For fibers containing at least one extreme gap (*E_i_* > 0), conditional severity was defined as

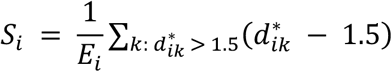

*S_i_* therefore measures the mean exceedance above the threshold rather than the absolute normalized gap size. Severity was left undefined for fibers without an extreme gap, because assigning a value of zero would conflate nonoccurrence with low conditional magnitude.

Age-adjusted genotype differences in extreme-gap frequency were estimated with a grouped binomial model containing categorical age and genotype. Conditional severity was modeled among fibers with *E_i_* > 0 using a Gamma generalized linear model with a log link and the same additive fixed effects. Genotype-specific estimates were standardized across development by predicting each outcome at all six ages and taking the equally weighted mean of the six age-specific predictions. This prevented ages with larger numbers of sampled fibers or mice from dominating the pooled genotype comparison.

Confidence intervals for standardized estimates and genotype differences were obtained from 10,000 stratified mouse-cluster bootstrap samples. Mice were resampled with replacement within each age-by-genotype stratum, both models were refitted in every bootstrap sample, and standardized predictions were recalculated. The 2.5th and 97.5th percentiles of the bootstrap distributions defined the 95% confidence intervals.

The association between fiber-level extreme-gap frequency and conditional severity was summarized with Spearman rank correlation among fibers containing at least one extreme gap. Correlations were estimated for the pooled dataset and separately within Control and Δ2w fibers. To estimate the association independent of the overall genotype difference, frequency and severity were rank transformed, each rank variable was residualized against genotype, and the residuals were correlated. Confidence intervals were obtained from 10,000 mouse-cluster bootstrap samples stratified by age and genotype. The conditional correlation analysis included 553 fibers (448 Control and 105 Δ2w) from 40 mice.

### Fiber-normalized one-dimensional pair-correlation analysis

The one-dimensional pair-correlation function was used to identify the relative distance scales at which myonuclear pairs were depleted or enriched. For fiber *i*, containing *N_i_* nuclei within a measured longitudinal segment of length *L_i_*, the even-spacing distance was defined as

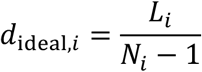

The separation between nuclei *k* and *l*, ℎ*_ikl_* = |*s_ik_* − *s_il_*|, was expressed on a fiber-normalized scale as

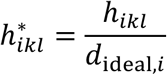

such that ℎ^∗^ = 1 corresponds to the separation obtained if the observed nuclei were distributed evenly across the analyzed segment. The measured fiber segment length *L_i_* was used as the fixed observation-window length for both distance normalization and edge correction. The corresponding normalized window length was

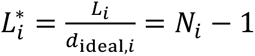

Fibers in which the projected nuclear span exceeded the measured segment length were excluded rather than enlarging the observation window on the basis of the observed nuclear positions. Forty-seven fibers met this criterion, leaving 982 fibers–482 Control and 500 Δ2w fibers–from 42 mice. At least three nuclei were required for inclusion.

The normalized pair-correlation function was evaluated in 40 equally spaced bins over 0 ≤ *r*^∗^ ≤ 4. For a bin *B_b_* with attainable width 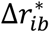, the per-fiber estimator was

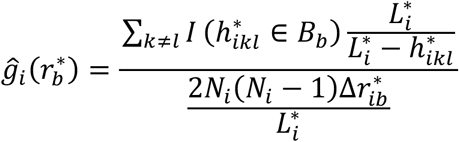

Thus, each ordered nuclear pair was weighted by the one-dimensional translation correction 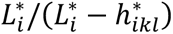 equivalent to *L_i_*/(*L_i_* − ℎ*_ikl_*) in physical units. The denominator is the conditional fixed-*N* CSR expectation after translation weighting. Consequently, 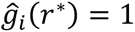 in expectation for independently and uniformly distributed nuclei within the fixed fiber window, whereas values below or above 1 indicate depletion or enrichment of nuclear pairs at that relative separation. Bins beyond the maximum attainable separation for an individual fiber were treated as missing.

Fiber-specific curves were first averaged across fibers from the same mouse. Age- and genotype-specific curves were then calculated as equal-weight means of the mouse-level curves. Pointwise 95% confidence intervals were obtained from 10,000 mouse-cluster bootstrap samples within each age-by-genotype stratum. In each bootstrap sample, mice were resampled with replacement and their complete mouse-level curves were retained. Enrichment around the expected even-spacing distance was summarized by the mean of *g*(*r*^∗^) over 0.85 ≤ *r*^∗^ ≤ 1.15. The binned curve was integrated over this exact interval, with boundary bins weighted according to their overlap with the interval. Age-specific Δ2w–Control contrasts and 95% confidence intervals were calculated using the same mouse-cluster bootstrap procedure. No pointwise hypothesis tests were performed across the full curve.

### Surface coordinate system and periodic Voronoi partitioning

Surface analyses used the same PCA-derived longitudinal axis as the one-dimensional analyses. Two orthogonal transverse vectors were obtained from the second and third principal components and re-orthogonalized numerically. For nucleus *k* in fiber *i*, the angular coordinate around the fiber axis was

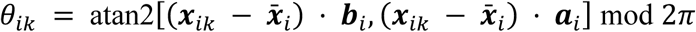

For surface analysis, each reconstructed fiber segment was represented by an equivalent cylindrical surface with the measured segment, *L_i_*, and surface area, *A*_surface,i._ The corresponding circumference was

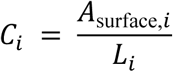

and the angular coordinate was converted to circumferential distance as

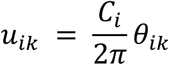

The resulting unwrapped surface window had dimensions *L_i_* × *C_i_* and therefore preserved the measured fiber surface area exactly. Let span*_i_* = max(*s_ik_*) − min(*s_ik_*). When span*_i_* ≤ *L_i_*, the portion of the measured segment length not occupied by the observed nuclear span was allocated symmetrically beyond the two outermost nuclei:

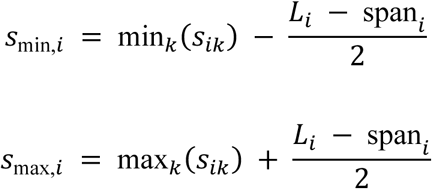

Fibers containing fewer than six nuclei were excluded. Fibers in which the observed nuclear span exceeded the measured segment length were also excluded rather than enlarging the analysis window. Of 1,015 otherwise eligible fibers, 47 met this criterion, leaving 968 fibers from 42 mice for surface analysis.

Periodic boundary conditions were imposed circumferentially by replicating each nuclear coordinate at *u_ik_* − *C_i_*, *u_ik_*, and *u_ik_* + *C_i_*. A planar Voronoi tessellation was then calculated. Unbounded regions were reconstructed as finite polygons, and all polygons were clipped to the fiber-specific rectangle [*s_min,i_*, *s_max,i_*] × [0, *C_i_*]. For each nucleus, the clipped fragments belonging to its three periodic copies were combined to recover the complete Voronoi cell within the central surface window. Numerical quality control confirmed that the summed Voronoi cell areas equaled the analyzed surface area, with only negligible numerical rounding error.

Surface partitioning heterogeneity was quantified as the coefficient of variation of the *N_i_* clipped Voronoi cell areas,

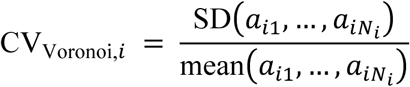

where *a_ik_* is the Voronoi cell area associated with nucleus *k* and the sample standard deviation used *N_i_* − 1 degrees of freedom. Lower Voronoi CV indicates more uniform partitioning of the analyzed fiber surface among the myonuclei.

### Nested spatial null models and dimensional decomposition

Two hierarchically related null models were used to separate the contribution of longitudinal positioning from additional circumferential organization. Both models preserved the observed nuclear number and the fiber-specific surface geometry. The fixed-s null additionally preserved every observed longitudinal coordinate while randomizing only the circumferential coordinate:

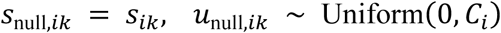

The fixed-s model therefore represents the surface organization expected if the measured longitudinal arrangement were retained but circumferential positions contained no additional structure. In the full 2D null model, both coordinates were randomized within the same fiber-specific surface window:

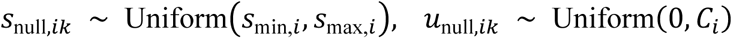

The full 2D model represents complete spatial randomness conditional on nuclear number and the measured analysis geometry. Each null model was simulated 1,000 times per fiber. Voronoi CV was recalculated for every realization using the same periodic replication, finite-region reconstruction, window clipping, and area calculation applied to the observed coordinates. All included fibers yielded 1,000 valid realizations for each null model. The mean null expectations are denoted *μ*_fixed-s,i_ and *μ*_full2D,i._

The reduction in Voronoi CV relative to complete two-dimensional randomness was decomposed additively into a longitudinal contribution and a residual circumferential contribution:

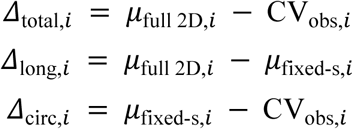

such that

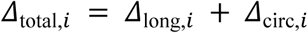

*Δ*_long_ quantifies the reduction in surface heterogeneity attributable to the observed longitudinal arrangement, whereas *Δ*_circ_ quantifies the additional reduction in heterogeneity remaining after longitudinal coordinates are held fixed.

For each mouse, *Δ*_total_, *Δ*_long_, and *Δ*_circ_ were first averaged across its analyzed fibers within each developmental age. Age- and genotype-specific component fractions were then calculated from mouse-balanced group means:

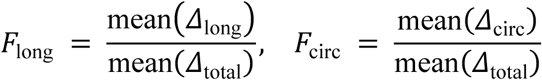

Uncertainty was estimated with 10,000 mouse-level bootstrap samples within each age-by-genotype stratum. Mice were sampled with replacement, and mouse-balanced component means and fractions were recalculated in each sample. 95% confidence intervals were defined by the 2.5th and 97.5th percentiles of the bootstrap distributions. This procedure ensured that animals, rather than the number of fibers sampled from each animal, determined the weighting of group-level estimates.

### Statistical analysis

The mouse was treated as the biological replicate for inferential analyses. Continuous fiber-level outcomes without a dedicated count or simulation-based model were analyzed with linear mixed-effects models containing genotype, categorical developmental age, and their interaction as fixed effects and a random intercept for mouse. The random intercept accounted for the non-independence of multiple fibers obtained from the same animal. Model-based genotype contrasts were estimated within each developmental age and reported with 95% confidence intervals. When six age-specific contrasts were evaluated for the same outcome, P values were adjusted using the Benjamini–Hochberg false-discovery-rate procedure.

The grouped binomial models, Gamma model, mouse-cluster bootstrap procedures, pair-correlation analysis, and Voronoi decomposition are described with their respective analyses above. Pair-correlation curves were summarized at the mouse level, and uncertainty was quantified using mouse-cluster bootstrap confidence intervals. The mean pair enrichment over 0.85 ≤ *r*^∗^ ≤ 1.15 was used as a prespecified scalar summary of organization around the expected even-spacing distance; no pointwise *P* values were calculated for the full curves. All inferential tests were two-sided, with α = 0.05. Effect estimates and confidence intervals were emphasized alongside P values.

Analyses were performed in Python 3.11 and R 4.5.3. PCA was implemented with scikit-learn 1.8.0, Voronoi tessellation with SciPy, and statistical modeling with statsmodels in Python and the base stats, lme4, and lmerTest packages in R. Data processing used pandas and tidyverse 2.0.0. Fixed random seeds were used for all simulation and bootstrap procedures to ensure reproducibility.

## Code availability

All analysis code is available from the corresponding author upon reasonable request.

## Author contributions

K.-A.H. conceived the study and wrote the original draft. Both authors performed the computational and statistical analyses, prepared the figures, developed the analytical framework, reviewed the manuscript, and approved the final version.

## Data availability

The datasets analyzed during the current study, including three-dimensional myonuclear coordinates and fiber-level geometric measurements, are available from the corresponding author upon reasonable request.

